# The SARS-CoV-2 spike reversibly samples an open-trimer conformation exposing novel epitopes

**DOI:** 10.1101/2021.07.11.451855

**Authors:** Shawn M. Costello, Sophie R. Shoemaker, Helen T. Hobbs, Annalee W. Nguyen, Ching-Lin Hsieh, Jennifer A. Maynard, Jason S. McLellan, John E. Pak, Susan Marqusee

## Abstract

Current COVID-19 vaccines and many clinical diagnostics are based on the structure and function of the SARS-CoV-2 spike ectodomain. Using hydrogen deuterium exchange mass spectrometry, we have uncovered that, in addition to the prefusion structure determined by cryo-EM, this protein adopts an alternative conformation that interconverts slowly with the canonical prefusion structure. This new conformation—an open trimer— contains easily accessible RBDs. It exposes the conserved trimer interface buried in the prefusion conformation, thus exposing potential epitopes for pan-coronavirus antibody and ligand recognition. The population of this state and kinetics of interconversion are modulated by temperature, receptor binding, antibody binding, and sequence variants observed in the natural population. Knowledge of the structure and populations of this conformation will help improve existing diagnostics, therapeutics, and vaccines.

**One Sentence Summary:** An alternative conformation of SARS-CoV-2 spike ectodomain modulated by temperature, binding, and sequence variants.

## Introduction

The spike protein from SARS-CoV-2 (also referred to as the S-protein) is the primary target for current vaccines against COVID-19 and the focus of many therapeutic efforts (*1*–*4*). This large heavily glycosylated trimeric protein is responsible for cell entry via recognition of the host receptor angiotensin-converting enzyme 2 (ACE2) and membrane fusion (*5*–*7*). It is also the principal antigenic determinant of neutralizing antibodies (*8*). Shortly after release of the viral genome sequence, a version of the spike ectodomain (termed S-2P) was designed to stabilize the prefusion conformation, and the structure was determined by cryo-electron microscopy (cryo-EM) (*9*, *10*).

This S-2P ectodomain comprises the first ~1200 residues of the spike protein (Fig. 1A) with two proline substitutions in the S2 domain designed to stabilize the prefusion conformation, mutations that abolish the furin-cleavage site, and the addition of a C-terminal trimerization motif (*9*). This mimic, its structure, and others that followed, have been widely used for vaccine development and interpretation of many structure/function and epidemiological studies. To date there are more than 250 structures of SARS-CoV-2 spike ectodomains in the Protein Data Bank (*11*).

**Figure 1.**
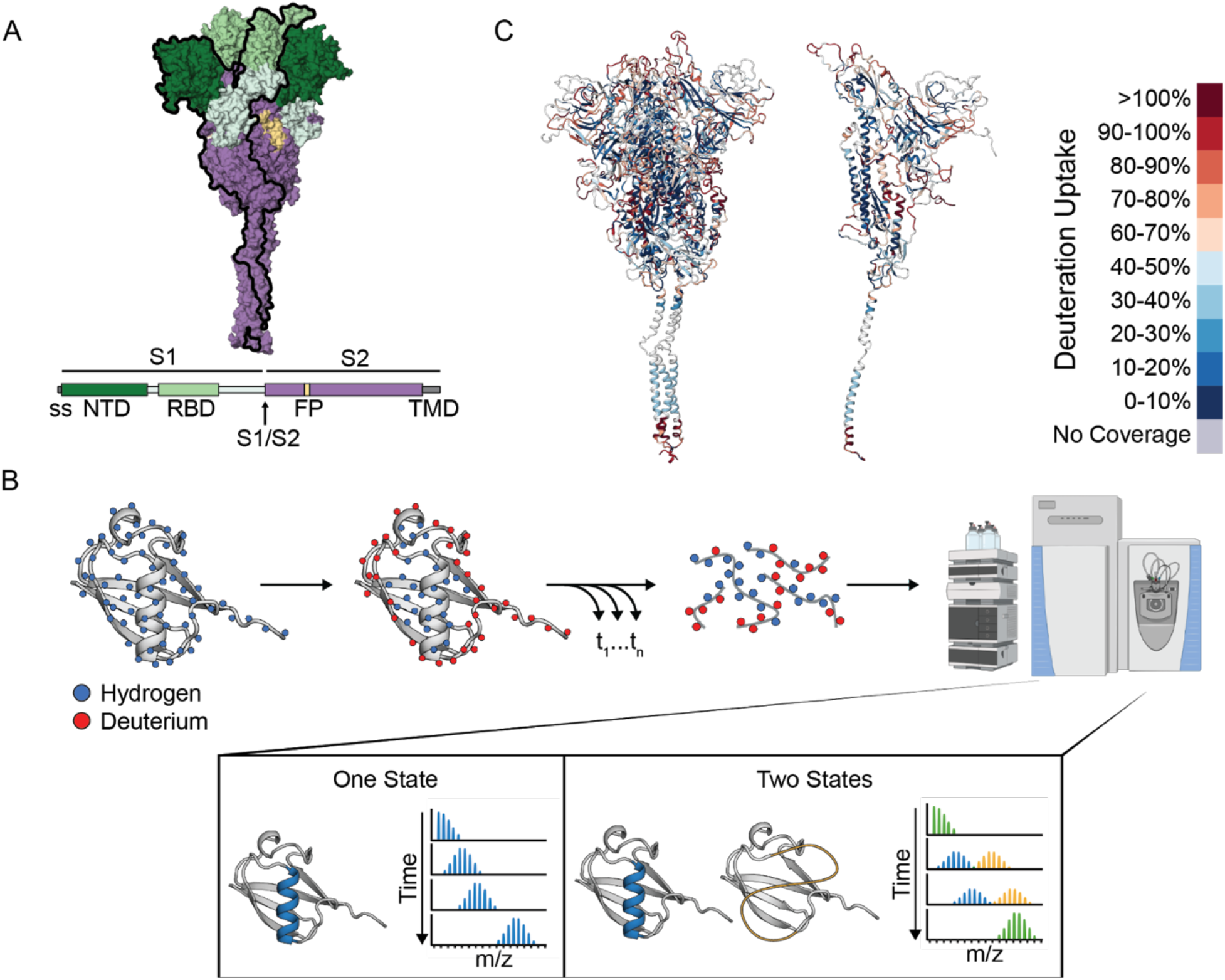
Hydrogen-Deuterium Exchange monitored by mass spectrometry (HDX-MS) on the SARS-CoV-2 Spike ectodomain. **(A)** Schematic of the prefusion-stabilized SARS-CoV-2 spike protein and a model of the trimeric prefusion conformation (24). **(B)** Schematic of HDX-MS experiment and the resulting mass distributions for a peptide that exists in either one (left) or two (right) separable conformations. In order for the two conformations to result in a bimodal mass distribution, they must not interconvert during the timescale of the HDX experiment (hours). Rapid interconversion would result in a single mass distribution with the ensemble averaged mass profile. **(C)** Schematic representation of the deuterium uptake across the entire spike protein displayed on the full trimer (left) or a single protomer (right) after 1 minute of exchange.

These structural studies together with other functional studies demonstrate that, like all class 1 viral fusion proteins, the spike protein is dynamic, sampling several different conformations during its functional lifecycle (*12*, *13*). The three individual receptor-binding domains (RBDs) sample an ‘up’ state and a ‘down’ state; the up state exposes the ACE2-binding motif and therefore is required for infectivity (*7*, *10*, *14*, *15*). After receptor binding and cleavage between the S1 and S2 domains, the protein undergoes a major refolding event to allow fusion and adopts the stable post-fusion conformation (*6*, *7*, *16*–*18*).

Despite the wealth of structural information, there are very few experimental studies on the dynamics within the prefusion state. The noted RBD up/down conformational transition has been monitored on the membrane via single molecule FRET and occurs on the order of seconds (*19*). Large computational resources have been devoted to molecular simulations of the spike protein revealing a dynamic pre-fusion state with a range of accessible conformations including the potential of a further opening of the RBD and N-terminal domain (NTD) away from the trimer interface (*20*, *21*). Experimentally, the conformational landscape of spike has not been well interrogated and the effects of perturbations, such as ligand binding (both receptor and antibodies) or amino acid substitutions found in emerging variants of concern are unknown.

For these reasons, we turned to hydrogen deuterium exchange monitored by mass spectrometry (HDX-MS) to probe the energy landscape of the soluble spike prefusion ectodomain as well as the effects of ligand binding and sequence variation. We uncovered a stable alternative conformation that interconverts slowly with the canonical prefusion structure. This conformation is an open trimer, with easily accessible RBDs. It exposes the S2 trimer interface, providing new epitopes in a highly conserved region of the protein.

## Results

### HDX-MS on Spike 2P (S-2P)

HDX-MS offers an ideal complement to the ever-growing number of structural studies on the SARS-CoV-2 spike protein, providing information on its conformational ensemble and dynamics. HDX-MS monitors the time course of exchange of amide hydrogens on the peptide backbone with the hydrogens in the solvent (see Fig. 1B for description). An individual amide’s ability to undergo exchange is directly related to its structure and stability (*22*, *23*).

We first followed the time course of hydrogen exchange on the entire S-2P ectodomain, over a period of 15 seconds to 4 hours (see Materials and Methods). Using a combination of porcine pepsin and aspergillopepsin digestion, we obtained 85% peptide coverage allowing us to interrogate the dynamics of the entire protein (800 peptides, which include 8 of the 22 glycosylation sites, average redundancy of 8.6) (fig. S1A). Notably, we have coverage in areas not resolved in the cryo-EM structure, including loops in the N-terminal domain (NTD) and RBD that have been found to be recognized by antibodies, loops in the S2 region that include the protease cleavage sites, and C-terminal residues after residue 1145 which includes the second heptad repeat (HR2). Based on control experiments using deuterated protein, our HDX protocol results in an average back exchange of 22% (fig S1B).

The vast majority of peptides show a classic single isotopic envelope whose centroid increases in mass as deuterons are added over time (Fig. 1B). A small minority of the peptides, however, show bimodal behavior—with two isotopic envelopes both increasing in mass over time: one less-exchanged distribution and a second more-exchanged distribution (these peptides are described in detail below). The HDX profile of all the peptides, with the exception of the more-exchanged distributions in the bimodal peptides, is consistent with the known prefusion conformation (Fig. 1C, fig. S2): secondary structure and buried elements within the trimer exchange slower than exposed loops. We also observe protection for residues 1140–1197, which includes HR2, a region not defined in single-particle cryo-EM structures, supporting the predicted helical structure of this region (*24*) and the relative rigidity of the stalk observed by cryo-electron tomography (cryo-ET) (*25*).

### Identification of an alternative conformation

Bimodal mass envelopes can indicate the presence of two different conformations that interconvert slowly on the timescale of our hydrogen exchange experiment: one where the amides are more accessible to exchange compared to the other. However, it can also be a result of the kinetics of the hydrogen exchange process itself, so-called EX1 exchange (when the rates of hydrogen bond closing are much slower than the intrinsic chemistry of the exchange process). In this rare scenario, the heavier mass distribution will increase in intensity at the expense of the lighter one over the observed time period. This is not what we observe for the spike protein: the bimodal mass distributions retain their relative intensities, increasing in average mass over time (fig. S3). The relative population of each state is the same for every bimodal peptide under any given condition. Thus, these bimodal peptides do indeed reflect two different conformations; they report on the regions of the protein that show differences in hydrogen exchange in each conformation.

The bimodal peptides we observe are predominantly in the most conserved region of the protein—the S2 region (*26*) (Fig. 2A). When mapped onto the canonical prefusion conformation, many come from the helices at the trimer interface (residues 962–1024, 1146–1166, 1187–1196), suggesting that these helices are either less stable or more solvent exposed in this newly identified conformation. We also observe bimodal peptides in other areas of the inter-protomer interface, such as residues 870–916 in S2 and residues 553–574 and 662–673 in S1, again suggesting a loss of trimer contacts. Finally, we see bimodal peptides in two regions that do not form interprotomer contacts (residues 291–305, 626–636); instead, these residues form the interface between the NTD and second S1 subdomain (SD2), suggesting that this subdomain interface is also lost. All the other peptides (the majority) fit to a classic unimodal distribution in the mass spectrum suggesting they behave similarly in both conformations. Previous HDX studies involving the spike protein did not note this behavior (*27*, *28*) (Supplementary Text).

**Figure 2.**
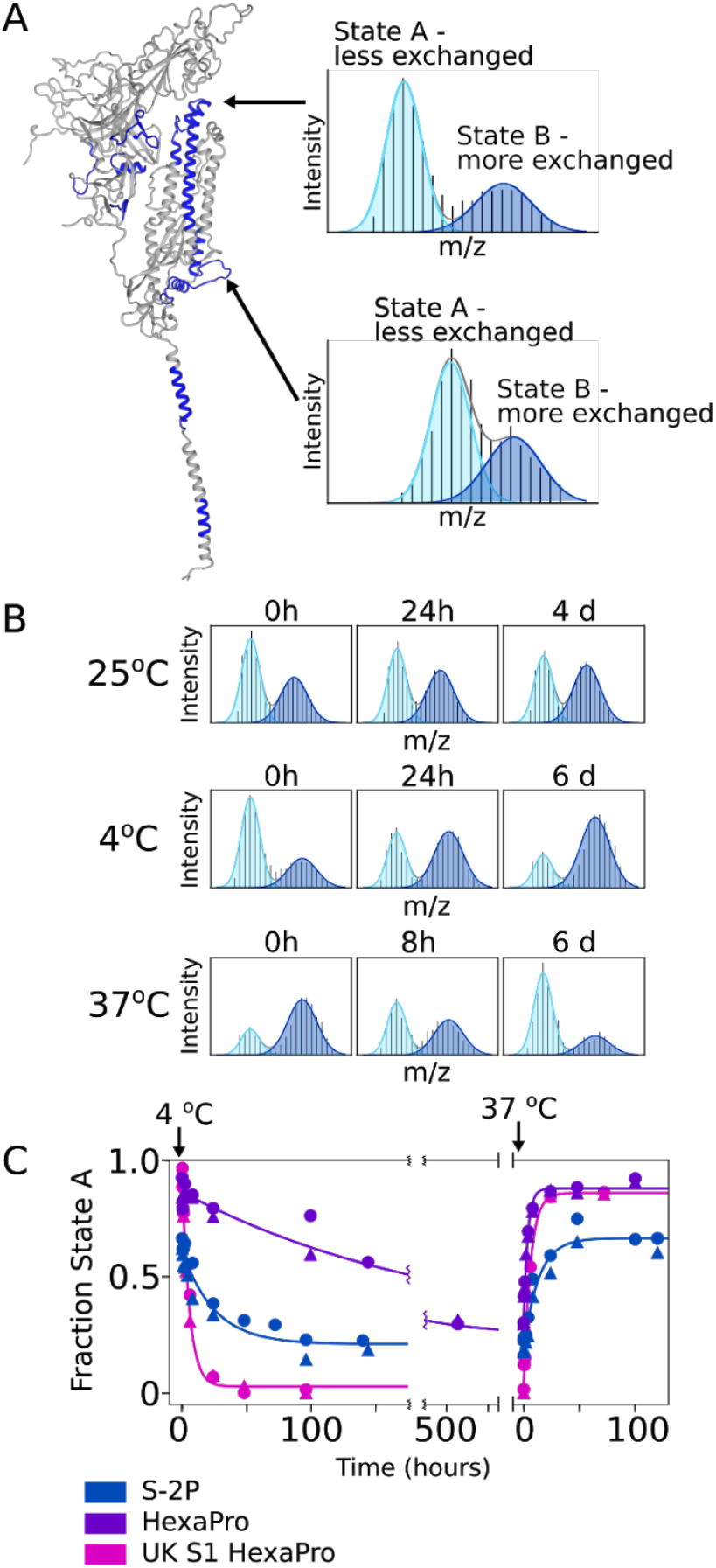
The Spike ectodomain reversibly samples two conformations. **(A)** Left: SARS-CoV-2 spike monomer with all regions that have peptides showing bimodal mass distributions colored in blue. Right: Example mass spectra from two peptides (top: residues 982–1001, bottom: residues 878–903) with overlaid fitted gaussian distributions that describe each protein conformation in blue (light blue: less exchanged A state, dark blue: the more exchanged B state). **(B)** Conformational preference for the S-2P spike construct at 25 °C, 4 °C and 37 °C. At 25 °C S-2P converts from primarily state A to ~50:50 A:B after 4 days. At 4 °C, S-2P prefers state B while at 37 °C, S-2P prefers state A. **(C**) The kinetics of interconversion between the A and B states for different of spike variants. Starting from an initial prefusion conformation (state A, 37 °C), samples were rapidly transferred to 4 °C and assayed for conversion to state B over time. To estimate fraction state A, peptides from two different regions (residues 982–1001 (circles) and residues 878–903 (triangles)) were fit to two gaussians. Data from both regions were used to determine the rate of interconversion.

### Interconversion between the two conformations

These data suggest a model where the spike protein populates two conformations within the prefusion state—the classical prefusion structure seen in cryo-EM (herein referred to as state A)—and one where each domain has a similar protomer topology and a more flexible or exposed open-trimer interface (herein referred to as state B). The above data do not, however, provide evidence that these states interconvert; any potential interconversion must be slower than the four-hour hydrogen-exchange experiment. Since the transition of the RBD between the up and down conformation occurs on the order of seconds, this conformational heterogeneity is not the source of the bimodal distributions and the observed hydrogen exchange reports on the weighted average of the two RBD conformations. There are several irreversible situations that could account for conformational heterogeneity such as differences in glycosylation, proteolytic degradation, irreversible misfolding, or aggregation. To rule these out, we tested whether the two conformations interconvert reversibly. We used the bimodal peptides to quantify the population of each conformation under differing conditions (such as temperature, time, ligand, etc.). Under each condition, we carried out a one-minute pulse of hydrogen exchange and integrated the area under the two mass envelopes for a single bimodal peptide to ascertain the fraction of each conformation under that condition or moment in time (see Materials and Methods). For every condition tested, irrespective of the A:B ratio, all of the peptides examined resulted in the same fractional population for each conformation, indicating that all of these data can be best described as a variable mixture of just two conformations: the canonical prefusion conformation and an unexpected alternative conformation.

Long-term incubation (four days, 25 °C, pH 7.4) demonstrated a slow shift in population from a majority in the canonical prefusion state (state A) to a majority in the alternative conformation (state B) (Fig. 2B). Thus, the prefusion state can transform into the alternative state and the bimodal behavior cannot be due to sample heterogeneity such as differential glycosylation. This observed A ⟶ B conversion, however, does not rule out an irreversible process such as degradation or misfolding.

Postulating that the bimodal peaks represent a reversible structural transition, we used temperature to perturb the system and investigate the ability to interconvert. Indeed, the conformations do interconvert reversibly, with a preference for B at 4 °C and A at 37 °C (Fig. 2B). The observed kinetics of interconversion are extremely slow: A ⟶ B t_1/2_ of ~17 hours at 4 °C and, when that same sample is moved to ~37 °C, B ⟶ A t_1/2_ of ~ 9 hours (Fig. 2C, Table 1). Notably the final distribution at either temperature shows an observable population of both states, indicating a very small energetic difference between the two conformations.

**Table 1.** Rates of interconversion between the A (prefusion) and B (open trimer) conformations of Spike ectodomains. *k_observed_* is the observed rate of change in the population of the A state after a temperature jump. This relaxation rate is the sum of the forward and reverse rates, which is dominated by the major conformational change (A⟶B at 4 °C, 10 °C and B⟶A at 37°C). t_1/2_ is the half time for that same rate, ln2/*k_obs_*).

| Temperature | Protein | $k_{observed}$<br>(hr<sup>-1</sup>) | $t_{1/2}$<br>(hours) |
| --- | --- | --- | --- |
| 37 °C→4 °C | S-2P | 0.04 | 17 |
|  | HexaPro | 0.005 | 143 |
|  | UK S1 HexaPro | 0.2 | 4 |
|  | S-2P + 3A3 | 0.1 | 5 |
| 37 °C→10 °C | S-2P | 0.05 | 14 |
|  | HexaPro | 0.004 | 171 |
|  | UK S1 HexaPro | 0.1, 0.2 (*) | 3, 5 (*) |
| 4 °C→ 37 °C | S-2P | 0.08 | 9 |
|  | HexaPro | 0.3 | 2 |
|  | UK S1 HexaPro | 0.2 | 4 |

(\*) Time course was monitored twice, and the results of each fit are reported.

The prefusion conformation of spike has already been noted to be temperature dependent. Cryo-EM experiments of spike incubated at 4 °C for 5 to 7 days showed less than 10% of the definable prefusion spike particles seen on grids of freshly prepared spike. Incubating spike at 37 °C for three hours after storage at 4 °C recovered particle density to the level seen using freshly prepared protein (*29*). Failure to detect particles also correlated with a loss in recognition by an antibody known to recognize quaternary structure. These studies are consistent with our findings—long-term incubation of spike at 37 °C biases to the prefusion conformation while long-term incubation at 4 °C prefers an expanded conformation, which is apparently not well visualized on cryo-EM grids.

### Effect of sequence changes—HexaPro

The small energetic difference between these states indicates that small changes in sequence may affect the relative populations and/or rates of interconversion between them. Indeed, the S-2P variant was designed to stabilize the pre-fusion conformation avoiding spontaneous conversion to the post-fusion form. This S-2P construct is the basis for most currently employed vaccines. Recently, a new version was constructed, termed HexaPro or S-6P, which contains four additional proline mutations designed to increase the apparent stability of the pre-fusion state and improve cellular expression (*30*).

Using the same HDX-MS process, HexaPro shows the same bimodal behavior, with the same regions reporting on the two conformations (see Fig. 2A, fig. S3B). At 4 °C, HexaPro, like S-2P, converts to state B, but with slower kinetics (t_1/2_ of ~6 days). At 37 °C HexaPro shifts back to state A with a t_1/2_ of ~2 hours (Fig. 2C). In sum, as expected based on the design criteria, HexaPro does result in a bias towards the prefusion conformation; however, both states are populated under all conditions, again consistent with two low energy conformations. Importantly, these changes demonstrate how a small number of mutations can perturb and modulate the conformational landscape of spike, suggesting that the evolving sequence variants may show differences in this conformational exchange (see below).

### Effects of sequence changes—an interprotomer disulfide bond

To further probe the structural features of the B conformation, we turned to a variant of HexaPro engineered to contain a disulfide bond. This variant trimer contains three disulfide bonds (S383C/D985C) that reach across protomers and lock the RBDs in the down state (*31*). When probed by HDX-MS we find that this disulfide-locked variant remains completely in the A state and does not show any observable population of the B state, even after O/N incubation at 25 °C. These data are consistent with a model where formation of the B state requires opening of the inter-protomer (trimer) interface and exposure of the RBDs, which would be prohibited by the interprotomer crosslinks.

### Effects of sequence changes—UK variant

Increasingly infectious SARS-CoV-2 variants of concern are being discovered throughout the global population on a regular basis. Most of these include mutations in the spike protein, primarily in the S1 domain; some reside in the ACE2-interaction surface, while others do not. Therefore, we asked if these mutations can influence the biases and kinetics of interconversion of the A and B conformation. We monitored the A/B conversion for a variant of HexaPro that includes five S1 mutations in the B.1.1.7 variant and none in the S2, (Δ69–70 (NTD), Δ144 (NTD), N501Y(RBD), A570D (SD1), P681H (SD2)), termed UK S1 HexaPro. Indeed, UK S1 HexaPro shows notable differences in both the relative preference for state B and the kinetics of interconversion. At 4 °C, UK S1 HexaPro converts to state B nearly 20 times faster than HexaPro (Fig. 2C, Table 1). Furthermore, UK S1 HexaPro shows no detectable prefusion conformer at 4 °C, while HexaPro shows at least 30% even after several weeks at 4 °C (fig S3c). At 37 °C, the kinetics and equilibrium distribution appear nearly identical between the two. All of the mutations are at solvent-exposed residues, except residue 570, which contacts the S2 subunit and resides in a region with observed bimodal behavior. Thus, despite their location in the S1 subunit and not at the core trimer interface, these specific B.1.1.7 mutations allosterically affect the interconversion of these two states.

### Effects of ACE2 binding

The primary function of the RBD is to recognize the host cell receptor ACE2. In the down conformation, the RBD is occluded from binding to ACE2, and in the up conformation its accessible. The entire trimer can exist with zero, one, two, or all three RBDs in the up conformation (*7*, *15*). In the isolated RBD, the Receptor Binding Motif (RBM) should always be accessible for ACE2 binding. We used HDX to monitor the binding of the receptor, both in isolation and in the full-length spike (S-2P), using a soluble dimeric form of ACE2 (ACE2-Fc, herein referred to as ACE2). For isolated RBD (residues 319–541, see Materials and Methods), we obtained 141 peptides, including one glycosylated peptide spanning the N-glycosylation site at residue 343 (no peptides are observed for site 331), resulting in 82% sequence coverage with an average redundancy of 8 (fig. S1).

The effects of ACE2 binding are illustrated in Figure 3. In the presence of ACE2, the latter half of the RBM (residues 472–513), shows a notable decrease in hydrogen exchange upon binding ACE2 (Fig. 3B), consistent with the known ACE2/RBD binding interface (*32*, *33*). We also observe small, but significant, changes for other regions that are near the binding interface. Importantly, we see very similar changes in HDX rates in RBD for both the isolated domain and in the context of the spike ectodomain, suggesting that all three RBDs in full-length spike interact with ACE2 in our experiment and that both the A and the B state can productively bind ACE2, which for the prefusion (A) state requires that RBD transition to the up state.

**Figure 3:**
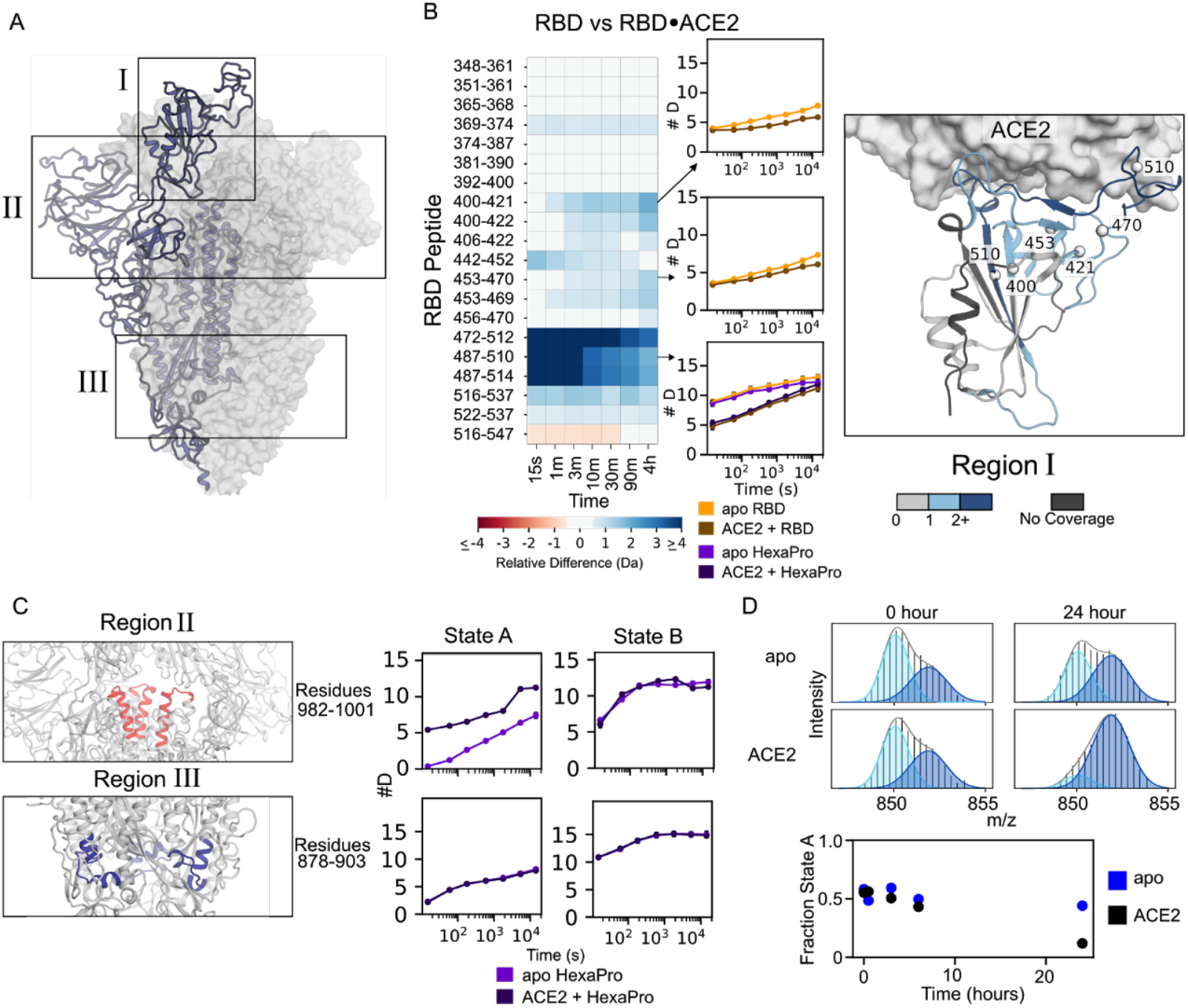
ACE2-binding effects are similar on isolated RBD and RBD in the context of the full-length ectodomain. **(A)** Diagram of the spike structure with regions of interest highlighted. **(B)** Left: Heatmap showing the difference in peptide deuteration in the presence and absence of ACE2 on the isolated RBD. Middle: The deuterium uptake plots are illustrated for three peptides of interest. The uptake plot for the RBM (residues 487– 510) is shown for both the isolated RBD and HexaPro. Right: Schematic representation of the heatmap data on the structure of the RBD•ACE2 complex (PDB 6M0J). The structure of the RBD is colored based on the maximum change shown in the heatmap for that residue in any peptide. **(C)** Changes to peptides from HexaPro upon binding of ACE2 outside of the RBD. When ACE2 binds the canonical prefusion structure, state A, peptide 982–1001 (Region II) becomes more solvent exposed and thus exchanges faster, but when ACE2 binds the open trimer (state B) it does not, presumably because it is already maximally exposed. For peptide 878–903 (Region III), there is no change in the exchange rate to either state A or state B indicating this region is not affected by ACE2 binding. In the schematic for region II, one NTD has been removed to visualize the peptide of interest. **(D).** Time course of interconversion in the presence of ACE2. Top: example spectra of S-2P peptide 878–902 with and without ACE2 before and after 24 hours of incubation at 25 °C. Bottom: time vs fraction state A for peptide 878–902 in S-2P with an without ACE2 over 24 hours. After 24 hours ACE2 bound S-2P prefers state B.

In the context of the spike trimer, we also observe notable changes outside of the RBD, particularly in state A, where a few peptides exchange more rapidly in the presence of ACE2 (in state B these peptides do not have any notable difference in the presence of ACE2) (Fig. 3C). These peptides are located on the top of S2 (residues 978–1001), a region known to become more exposed when RBD transitions from a down to an up conformation. Since ACE2 binding in the prefusion state requires the RBDs to be in the up conformation, this increased exchange reflects the known biases in the RBD conformation—a prefusion state whose RBDs are primarily in the down conformation and must transition to an up conformation to bind ACE2. We also see changes in the interconversion between state A and state B in the presence of ACE2, such that state B is more preferred (Figure 3D).

### The dynamics of the RBD are similar in isolation and the intact spike trimer

The isolated RBD has been used for many biochemical studies and is the main component of many clinical diagnostics. It is therefore important to ask whether there are large differences in the RBD when it is in isolation versus in the context of the spike trimer; our experiments allow us to directly compare the two. Very few peptides in the RBD show substantial changes in HDX behavior (fig. S4) and support the use of approaches such as deep sequence mutagenesis on the isolated RBD to gain information on the potential effects of variants, such as escape mutations (*34*).

We do, however, see some key differences in the RBD—mostly at the termini of the isolated domain and in the expected interactions with the rest of spike and across the protomer interface. The C-terminal region of the RBD (residues 516–537) is notably less protected in the isolated domain. This region is not part of the RBD globular domain and in full-length spike forms part of subdomain 1, so it is not surprising that there would be an increase in flexibility when isolated from the rest of this subdomain. Future studies with the isolated RBD may benefit from removal of both C-terminal and N-terminal (no peptide coverage observed for this region) regions, as they are likely disordered and may interfere with crystallization or lead to increases in aggregation.

### 3A3—an antibody that binds specifically to the B state

Recently, an antibody, 3A3, was developed that binds to MERS-CoV, SARS-CoV-1, and SARS-CoV-2, with an apparent epitope in a region where we observe bimodal behavior (residues ~980–1000) (*27*). This region, however, is inaccessible in the prefusion structure—it is buried in the prefusion structure when all RBDs are down, and highly occluded when the RBDs are up. Our HDX data indicate that this region is exposed in state B. To confirm the epitope, we repeated the HDX studies in the presence of 3A3; indeed, we see strong increased protection in the 978–1001 region. Moreover, this protection is directly associated with state B (Fig 4A). In state B, the epitope is now occluded from solvent and shows similar exchange in both the A and the B state. These data suggest a model where the 3A3 antibody binds uniquely to the B state.

**Figure 4:**
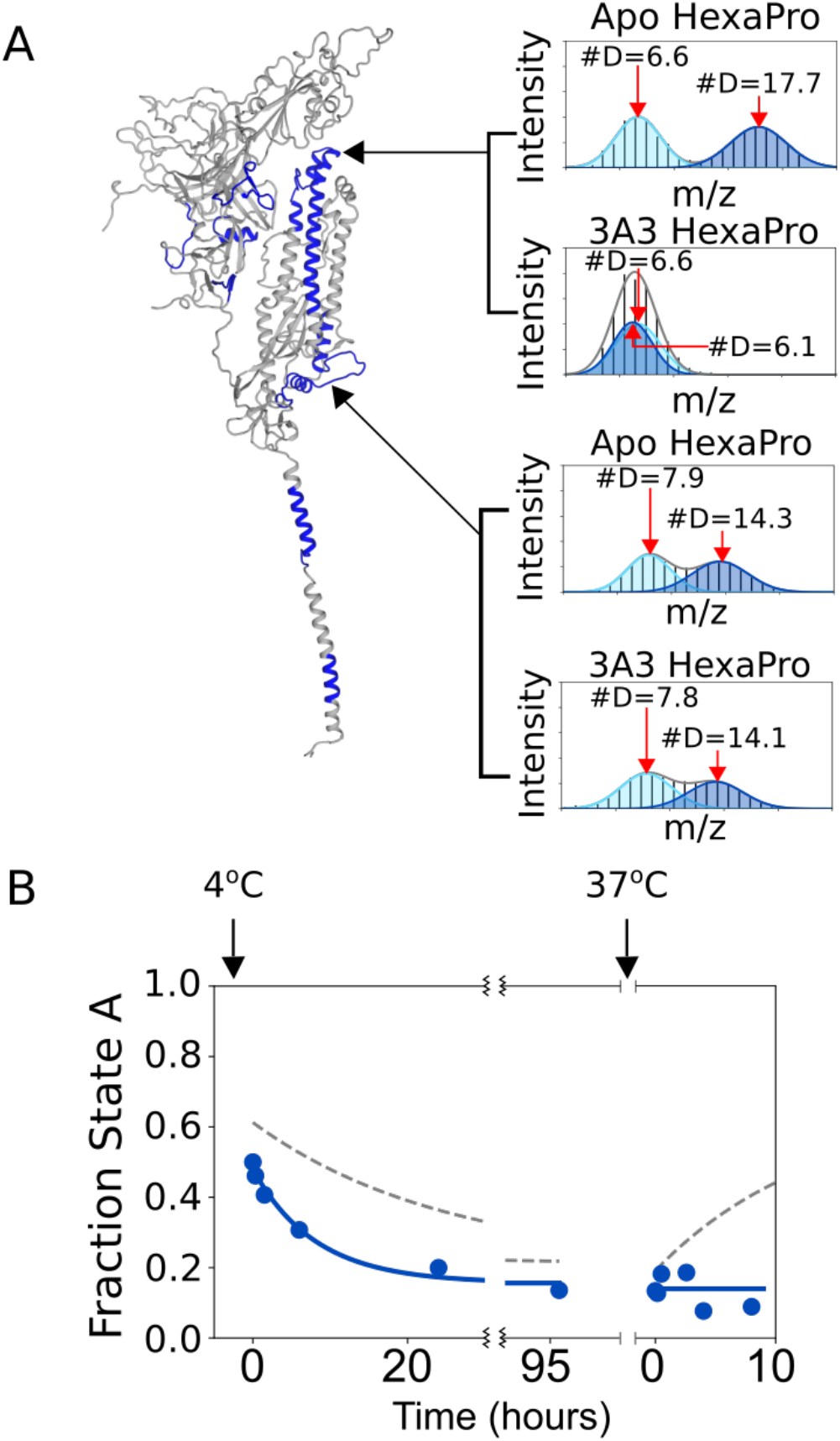
The antibody 3A3 binds to the region 982–1001 in state B. **(A)** Example mass spectra for HexaPro with and without 3A3 for two different peptides that have bimodal mass distributions. The bottom peptide shows no change in the presence of 3A3, which indicates that the amount of state A and state B has not significantly changed 13 minutes after adding 3A3. The top peptide, however, shows significant protection in the presence of 3A3, shifting the distribution belonging to state B to a deuteration amount indistinguishable from state A. **(B)** The kinetics of interconversion of S-2P in the presence of 3A3. The addition of 3A3 accelerated the rate of conversion to state B at 4 °C. The binding of 3A3 prevents the return to state A at 37 °C. Dotted lines indicate the conversion in the absence of 3A3.

To confirm this hypothesis, we looked at the effect of 3A3 binding on the temperature-induced conversion between A and B. 3A3 increases the rate of conversion from A to B at 4 °C, decreasing the t_1/2_ from ~17 hours to ~5 hours (Fig 4B, Table 1). This increase in the observed rate implies that 3A3 also affects the transition state for the conversion. Furthermore, returning the sample to 37 °C in the presence of 3A3 (state B saturated with 3A3) prohibits any transition back to the prefusion state, indicating that the binding of 3A3 prevents formation of the prefusion state most likely due to steric hindrance of the antibody being bound to the trimer interface. Since 3A3 binds wild-type and D614G spike when expressed on the surface of cells and neutralizes pseudovirus expressing these spikes (*27*), the data collectively suggest state B exposes broadly neutralization-sensitive epitopes that may be of interest for future therapeutics and vaccines.

### Structural model for state B

The above data allow us to create a structural model for state B (Fig. 5). The overall fold, or topology, of each domain is likely similar to the prefusion structure as, with the exception of the bimodal peptides, their hydrogen exchange patterns are similar. The bimodal peptides, which report on the two different conformations, cluster in the trimer interface, suggesting that this interface is more accessible to solvent in state B. State B is not a monomer. Size-exclusion chromatography and the hydrogen exchange data at the trimerization motif, confirm that both conformations are trimeric (fig. S5). In these soluble ectodomain constructs, the trimer is held together by the appended C-terminal trimerization domain, while in the full-length native spike trimer, the transmembrane helical segment likely serves this function. Therefore, state B is best modeled as an opened-up trimer with three protomers whose domains are structurally uncoupled. Our data do not let us address the relative orientation of the protomers within individual trimers; however, an ensemble of opened-up trimers with heterogeneous positioning of the protomers would best explain the lack of cryo-EM data. An opened-up class 1 viral fusion protein has been reported for respiratory syncytial virus (RSV) and visualized by a low resolution structure (*35*). This structural data from RSV and reports of an opening up of other viral fusion proteins (*36*–*38*) support our model of an ensemble of open-trimers with various degrees of openness.

**Figure 5.**
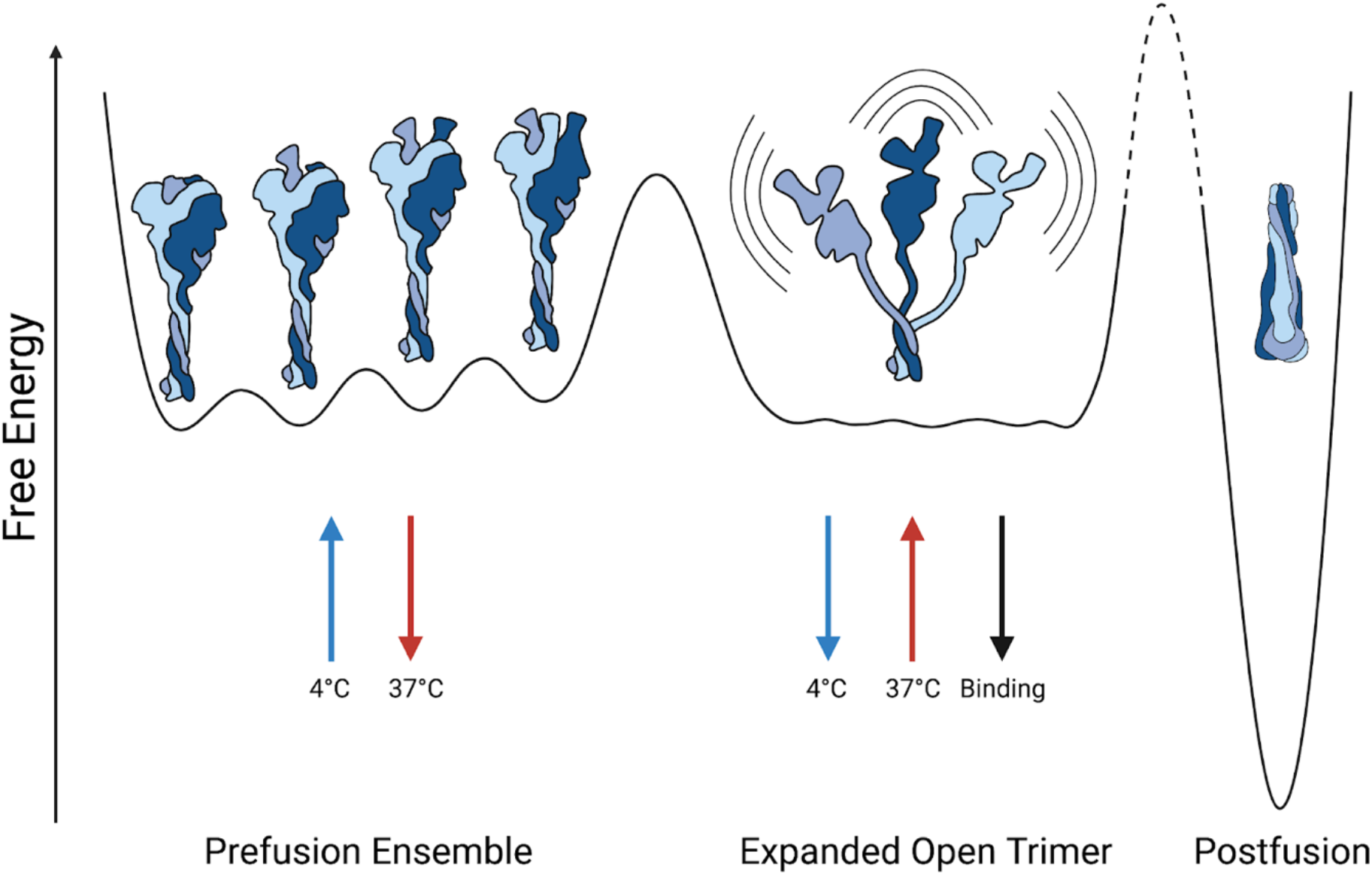
Schematic of the energy landscape for the SARS-CoV-2 Spike ectodomain. Reaction coordinate illustrating the different conformations accessible by Spike. Three different conformational states are depicted: the canonical prefusion ensemble, the expanded open trimer, and the postfusion conformation. The prefusion conformation contains all four RBD states (0,1,2, or 3 up). The relative energies and barrier heights as well as the placement of the open trimer along the reaction coordinate are drawn for illustration only.

A loss of interprotomer contacts in state B implies that the RBDs no longer contact adjacent protomers, and thus do occupy distinct ‘down’ and ‘up’ conformations; rather, they are likely always in a binding-competent state, perhaps even more accessible than the canonical ‘up’ state. This increased availability of the RBD may drive a preference for the B state in the presence of ACE2. Furthermore, in the prefusion conformation, having all three RBDs bound to ACE2 could lead to steric hindrances, but in state B, all three RBDs should be able to bind ACE2 with high affinity. Interestingly, introduction of variants of concern, such as in the UK HexaPro variant greatly increases the rate of conversion to state B, which may play a role in the noted increased infectivity.

Molecular dynamics has shown a smaller opening of the spike protein where an RBD and adjacent NTD twist and peel away from the center of the spike protein, revealing a cryptic epitope at the top of the S2 domain (*20*). This rapidly sampled conformation is not state B, as it does not involve the S2 trimer interface and the timescale of conversion to state B is unlikely to be sampled during a molecular dynamics simulation. This partial opening, however, could be on pathway to state B.

### A potential role for state B in spike function

The increase in the formation of state B upon binding ACE2 suggests that state B may be a functional intermediate. It may be an intermediate along the pathway to S1 shedding during the transition from the prefusion conformation to the postfusion conformation. This irreversible transition is not possible in the soluble ectodomain which removes the proteolytic cleavage site. If this is an on-pathway intermediate, antibodies or ligands that trap the protein in this state may block the protein along the pathway to fusion (*39*) those that act on the transition state and increase the rate of formation of the intermediate could promote the premature formation of the post-fusion conformation and thus aid in neutralization. If instead, formation of state B is off-pathway, antibodies or other ligands that favor state B would essentially trap the protein in an inactive conformation.

### The presence of this alternative conformation presents new druggable sites

The newly identified B state contains a large and unique solvent accessible surface area that is buried in the canonical prefusion conformation, thus exposing different epitopes for antibody and ligand recognition. Moreover, these regions arise from the most highly conserved part of the protein, the S2 trimer interface, and therefore present an ideal target for vaccines that would provide protection across coronaviruses (pan-coronavirus). In terms of therapeutics, ligands directed towards this region may also be broadly efficacious against variants of concern as well as other coronaviruses. Indeed, the antibody 3A3 represents one such potential therapeutic.

### The presence of state B will affect measured ligand-binding affinities

Finally, independent of whether this additional conformation is an on-pathway intermediate in the coronavirus functional lifecycle, it is ubiquitous among *in vitro* preparations of the spike protein. We see evidence of this conformation in every variant examined, excluding the disulfide-locked sample. Many biochemical and diagnostic assays use these isolated spike constructs, and many laboratories store this protein at 4 °C, where the alternative, expanded conformation is favored. Given that state A and B have differing affinities for the receptor and some antibodies, the temperature and time-dependent changes in the population of state B complicates quantitative analysis of binding affinities and needs to be further evaluated.

## Conclusion

In sum, we have found that the SARS-CoV-2 spike ectodomain reversibly samples an open-trimer conformation, allowing for the development of pan-coronavirus vaccines and therapeutics. This open trimer is folded and exposes a highly conserved region of the protein. It is similar in energy to the well-characterized prefusion conformation determined by cryo-EM. The fraction of spike in each conformer depends on temperature, ligands, and sequence. Mutations, receptor binding, antibodies, and temperature all affect the kinetics and energetics of this conformational change. Thus, quantitative measurements, such as *in vitro* binding assays, need to be re-evaluated for possible effects due to this mixed population. How easily this conformation is sampled in natural membrane-bound spike and its position in the viral lifecycle are still unknown; however, an antibody specific for this conformation can bind and neutralize in *in vitro* SARS-CoV-2 cell fusion and pseudovirus assays suggesting an important role. Determining which conformation elicits more robust protective immunity against both SARS-CoV-2 variants and other coronaviruses will be important for future vaccine development.

## Methods

### Protein Expression and Purification

SARS-CoV-2 Spike (2P) and RBD were expressed and purified from stably transformed Expi293 cells, following methods as described (*40*). HexaPro, HexaPro S383C/D985, and UK HexaPro were expressed and purified from transiently transfected ExpiCHO cells as described (*30*). ACE2-Fc was a gift from the Wells lab (*41*). 3A3 IgG was expressed and purified from ExpiCHO cells as described (*27*).

### Continuous Hydrogen exchange Labeling

For all continuous hydrogen exchange experiments, deuterated buffer was prepared by lyophilizing PBS (pH 7.4, Sigma-Aldrich P4417) and resuspending in D2O (Sigma-Aldrich 151882). To initiate the continuous labeling experiment, samples were diluted 10-fold (final spike trimer concentration of 0.167 μM) into temperature equilibrated deuterated PBS buffer (pH_read_ 7, pD 7.4). Samples were quenched, at the timepoints outlined below, by mixing 30 μL of the partially exchanged protein with 30 μL of 2x quench buffer (3.6 M GdmCl, 500 mM TCEP, 200 mM Glycine pH 2.4) on ice. Samples were incubated on ice for 1 minute to allow for partial unfolding to assist with proteolytic degradation and then flash frozen in liquid nitrogen and stored at −80°C.

For studies comparing HexaPro ± ACE2 the RBD in isolation vs in S-2P, purified spike (1.67 μM spike trimer or 5 μM RBD) was incubated in PBS at 25°C overnight (12-16 hours) before the initiation of hydrogen exchange. For experiments done in the presence of ACE2-Fc, the ligand was added during this incubation at a 1.25:1 molar ratio of ligand to spike monomer (6.25 μM ligand) to ensure saturation. Based on the reported affinity (K_D_ ~15 nM) for ACE2-Fc, fraction bound can be assumed to be greater than 97%. The hydrogen exchange time points for these experiments were 15 seconds, 60 seconds, 180 seconds, 600 seconds, 1800 seconds, 5400 seconds, and 14400 seconds.

For the comparison of HexaPro ± 3A3, HexaPro was incubated overnight at 37 °C (12-16 hours). After incubation the protein was moved to 25 °C and diluted to 1.67μM spike trimer. In the 3A3 bound condition, 6.25 μM antibody was added and allowed to bind for 10 minutes at 25 °C. Given the affinity of 3A3 for HexaPro (12 nM, fraction bound can be assumed to be greater than 97%. The quench time points for this experiment were 15 seconds, 180 seconds, 1800 seconds and 14400 seconds.

### Back exchange control preparation

S-2P was diluted to 1.67 μM trimer in PBS pH 7.4. To initiate hydrogen exchange, the sample was diluted 10-fold (final spike trimer concentration of 0.167 μM) into deuterated PBS buffer (pH_read_ 7, pD 7.4) that was supplemented with 3.6 M GdmCl, and then incubated at 37 °C. The addition of denaturant and increased temperature should both promote hydrogen exchange, by destabilizing folded structures and increasing the intrinsic rate of hydrogen exchange, respectively. Following two weeks of exchange, 30 μL of deuterated spike was mixed with 30 μL of 2X quench buffer lacking denaturant (500 mM TCEP, 200 mM Glycine pH 2.4) and kept on ice for one minute prior to flash freezing in liquid nitrogen and storage at −80 °C. The results of this control experiment were used to characterize the back exchange of the system and were not used to adjust deuteration values of continuous-labeling experiments.

### Incubation Kinetics and Pulse Labeling

For evaluating the temperature dependent kinetics of interconversion, frozen spike samples were thawed, diluted to 5 μM spike monomer, and incubated at 37°C overnight. Samples were then moved to a temperature-controlled chamber at 4°C and the population of each state was evaluated at the specified time points as described below. After the final 4 °C sample was taken (96–526 hours, depending on the spike construct), the sample was returned to a 37° heat block for further incubation and the population of each state was again evaluated at the specified time points as described below. To evaluate the relative population of the A and B conformer at each time point, 3 μL of spike sample was removed from the incubation tube and mixed with 27 μL of room temperature deuterated buffer. After a 1-minute labeling pulse, 30 μL of quench buffer kept on ice was mixed with the 30 μL of labeled protein. Quenched samples were kept on ice for 1 minute to allow for partial unfolding, and then flash frozen in liquid nitrogen.

For kinetics carried out in the presence of ACE2 or 3A3, after the initial 37 °C incubation the sample was brought to 25 °C and ligand was added (6.25 μM). To monitor the population of state A and B as a function of time at 25 °C in the presence of ACE2 the sample was kept in a temperature-controlled chamber at 25 °C and aliquots were removed for pulse labeling as described above at 0 hours, 30 minutes, 3 hours, 6 hours and 24 hours.

### Protease digestion and LC/MS

All samples were thawed immediately before injection into a cooled valve system (Trajan LEAP) coupled to a LC (Thermo UltiMate 3000). Sample time points were injected in random order. The temperature of the valve chamber, trap column, and analytical column were maintained at 2°C. The temperature of the protease column was maintained at 10°C. The quenched sample was subjected to inline digestion by two immobilized acid proteases in order, aspergillopepsin (Sigma-Aldrich P2143) and porcine pepsin (Sigma-Aldrich P6887) at a flow rate of 200 μL/min of buffer A (0.1 % formic acid). Protease columns were prepared in house by coupling protease to beads (Thermo Scientific POROS 20 Al aldehyde activated resin 1602906) and packed by hand into a column (2mm ID x 2 cm, IDEX C-130B). Following digestion, peptides were desalted for 4 minutes on a hand-packed trap column (Thermo Scientific POROS R2 reversed-phase resin 1112906, 1 mm ID x 2 cm, IDEX C-128). Peptides were then separated with a C8 analytical column (Thermo Scientific BioBasic-8 5 μm particle size 0.5 mm ID x 50 mm 72205-050565) and a gradient of 5-40% buffer B (100% Acetonitrile, 0.1% Formic Acid) at a flow rate of 40 μL/min over 14 minutes, and then of 40-90% buffer B over 30 seconds. The analytical and trap columns were then subjected to a sawtooth wash and equilibrated at 5% buffer B prior to the next injection. Protease columns were washed with two injections of 100 μL 1.6 M GdmCl, 0.1% formic acid prior to the next injection. Peptides were eluted directly into a Q Exactive Orbitrap Mass Spectrometer operating in positive mode (resolution 70000, AGC target 3e6, maximum IT 50 ms, scan range 300-1500 m/z). For each spike construct, a tandem mass spectrometry experiment was performed (Full MS settings the same as above, dd-MS^2^ settings as follows: resolution 17500, AGC target 2e5, maximum IT 100 ms, loop count 10, isolation window 2.0 m/z, NCE 28, charge state 1 and ≥7 excluded, dynamic exclusion of 15 seconds) on undeuterated samples.

### Peptide Identification

Byonic (Protein Metrics) was used to identify unmodified and glycosylated peptides in the tandem mass spectrometry data. The sequence of the expressed construct, including signal sequence and trimerization domain, was used as the search library. Sample digestion parameters were set to non-specific. Precursor mass tolerance and fragment mass tolerance was set to 6 and 10 ppm respectively. Variable N-linked glycosylation was allowed, with a library of 132 human N-glycans used in the search. No non-glycosylated peptides spanning any of the 22 known glycosylation sites in the spike sequence were ever observed, independent of the glycosylation search parameters. Peptide lists (sequence, charge state, and retention time) were exported from Byonic and imported into HDExaminer 3 (Sierra Analytics). When multiple peptide lists were obtained, all were imported and combined in HDExaminer 3.

### HDExaminer 3 Analysis

Peptide isotope distributions at each exchange time point were fit in HDExaminer 3. For glycosylated peptides, only the highest confidence modification was included in the mass spectra search and analysis. For unimodal peptides, deuteration levels were determined by subtracting mass centroids of deuterated peptides from undeuterated peptides. For bimodal peaks, extracted peptide isotope spectra were exported from HDExaminer 3 and analyzed separately (see below for details).

### Bimodal Fitting and Conformation Quantification

Peptide mass spectra for bimodal peptides were exported from HDExaminer 3.0. All quantitative analysis of the exported peptide mass spectra was performed using python scripts in Jupyter notebooks. After importing a peptide mass spectra, the m/z range containing all possible deuteration states of the selected peptide was isolated and the find_peaks method from the scipy.signal package was used to identify each isotope in the mass envelope and the height of each peak was used as its intensity. The area of the total mass envelope was normalized to account for run-to-run differences in intensity. The bimodal mass envelopes for all timepoints under the same condition were globally fit to a sum of two gaussians, keeping the center and width of each gaussian constant across all incubation time points. Fitting was done using the curve_fit function from the scipy.optimize package. After fitting, the area under each individual gaussian was determined to approximate the relative population of each state.

## Acknowledgements

We thank the entire Marqusee Lab for advice and critical reading of the manuscript, particularly Johanna Lindner for help initiating these experiments. We thank Peter S. Kim, Abigail Powell, Payton Weidenbacher, Natalia Freedland for advice and discussion; Jim Wells, Shion Lim and Irene Lui for discussion and providing ACE2-Fc and Aashish Manglik for advice and discussion. We thank Jeff Morrow and Sierra Analytics for making HDExaminer freely available for at-home use during shelter-in-place orders. SM is a Chan Zuckerberg Biohub Investigator. Some figures were created with the use of BioRender.com.

## Funding

National Institutes of Health grant R01GM050945 (SM), R01 AI127521(JSM), NIH AI122753 (JAM)

National Science Foundation grant MCB 1616591 (SM)

National Science Foundation GRFP (SMC)

Welch Foundation grant number F-0003-19620604

## Author Contribution

Conceptualization: SMC, SRS, HTH, SM

Methodology: SMC, SRS, HTH, JEP, SM

Investigation: SMC, SRS, HTH, JEP, AWN, CLH

Writing: SMC, SRS, HTH, AWN, CLH, JAM, JCM, JEP, SM

## Competing Interests

S.M.C., S.R.S. and S.M. are inventors on U.S. patent application no. 63/220,388, (“Methods related to an alternative conformation of the SARS-CoV-2 Spike Protein”). Y.H., A.W.N., C.-L.H., J.S.M. and J.A.M. are inventors on U.S. patent application no. 63/135,913 (“Cross-reactive antibodies recognizing the coronavirus spike S2 domain”). J.S.M. is an inventor on U.S. patent application no. 62/412,703 (“Prefusion Coronavirus Spike Proteins and Their Use”). C.-L.H., A.M.D., J.A.M., and J.S.M. are inventors on U.S. patent application no. 63/032,502 (“Engineered Coronavirus Spike (S) Protein and Methods of Use Thereof”).

## Supplementary Materials

Supplementary Text

Figs. S1 to S5

