## Supplementary Material for "The SARS-CoV-2 spike reversibly samples an open-trimer conformation exposing novel epitopes"

**Supplementary Text**

<u>Lack of observed bimodal peptides in previous SARS-CoV-2 spike HDX-MS studies</u>

Other groups have studied the spike protein by HDX-MS yet have not reported the presence of bimodal mass distributions for any peptides. We believe this discrepancy between our study and previous studies is largely due to differences in peptide coverage and back exchange. In Raghuvamsi et. al, many of the areas where we observed peptides with bimodal mass distributions in our data lacked peptide coverage in their study and more specifically lacked peptides long enough for obvious bimodal mass distributions. For one example, one of the bimodal peptides we often used for quantification of the two states reported in this paper was residues 878–902, this 24-residue peptide, with 21 exchangeable amides, is long enough to provide two very distinct mass distributions, one centered around approximately 9 added daltons and the other centered around 16 daltons. This 7 dalton separation is quite obvious. On the other hand, in Rahuvamsi et. al the longest peptide they saw in this region is reported to be 878–893, this peptide is only 15 residues with 13 exchangeable sites. We also have this peptide in our dataset, and it does appear bimodal, but the difference is less distinct, the lighter distribution is centered around 6.5 daltons while the heavier distribution is centered around 10.4 daltons. This 4 dalton difference is quite small and can make the bimodal mass distributions appear more like a skewed unimodal mass distribution. Furthermore, differences in back exchange and theoretical maximum deuteration could further obscure the two distributions. Our back exchange is estimated to be an average of 22% among peptides, Raghuvamsi et al estimate their back exchange to be an average of 34% back exchange. If we look at the previous example peptide, if Raghuvamsi et. al analyzed this peptide with 34% back exchange instead of 22% back exchange, they would see the lighter peak around 6 and the heavier peak around 9, further reducing the difference in center of the peaks to only 3 daltons, which would probably just appear as a single unimodal distribution. Finally, The construct in Raghuvamsi et. al is different from the ones we used in this study, notably it appears to not be glycosylated. While we have no evidence that glycosylation would or would not affect the bimodal mass distributions, glycosylation has been reported to affect different aspects of spike dynamics and thus could be a confounding variable.

Similarly in Huang et. al where they report their 3A3 antibody, they also do not report bimodal mass distributions for any peptides, even though the binding epitope of the antibody is a peptide we report with a bimodal mass distribution. As in Raghuvamsi et. al, we believe that the lack of coverage, lack of long peptides and differences in back exchange is the primary reason for this difference. Furthermore, the purpose of this preprint was not to analyze the HDX-MS behavior of the spike protein but was rather to map a binding epitope. Thus, it is reasonable to expect a thorough examination of every peptide in the protein was not part of the analysis and thus bimodal peptides could have been missed.

A

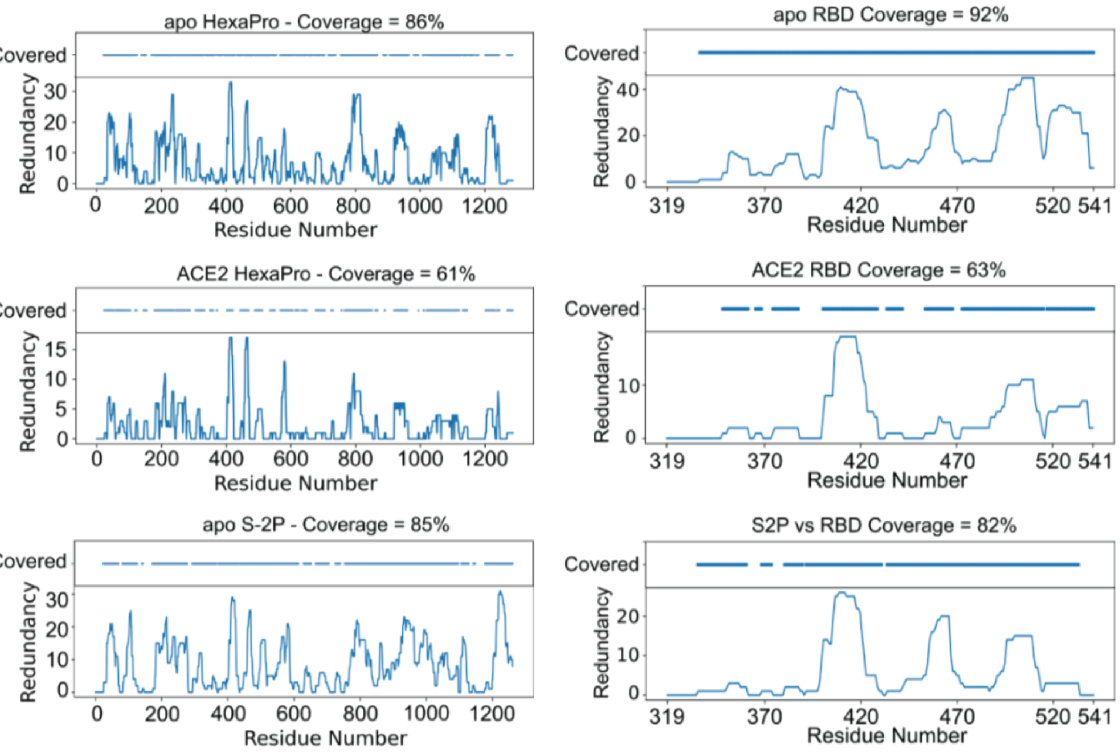


B

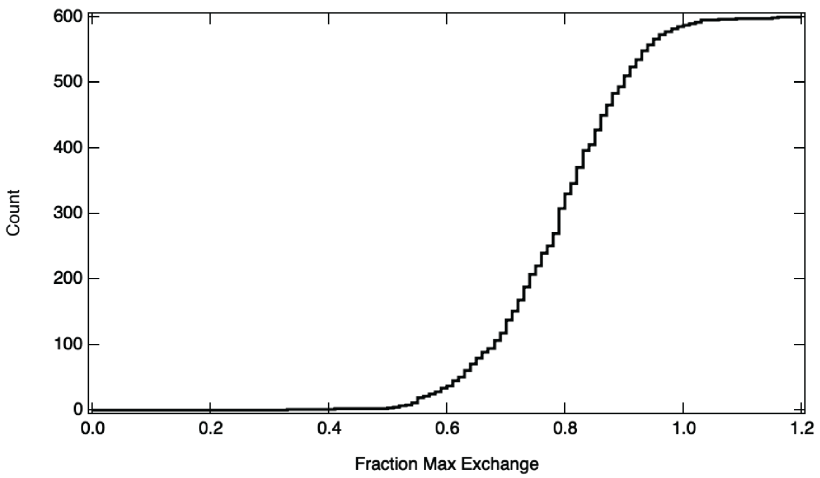


**Fig. S1. Coverage, redundancy, and back exchange results from HDX-MS experiments**
**(A)** Peptide coverage and redundancy at each residue for all HDX-MS experiments. **(B)** Back-Exchange Control: Cumulative histogram of the fractional deuterium maintained during workup of a fully deuterated sample. Fraction max exchange is corrected for the 90% $D_2O$ experimental conditions.

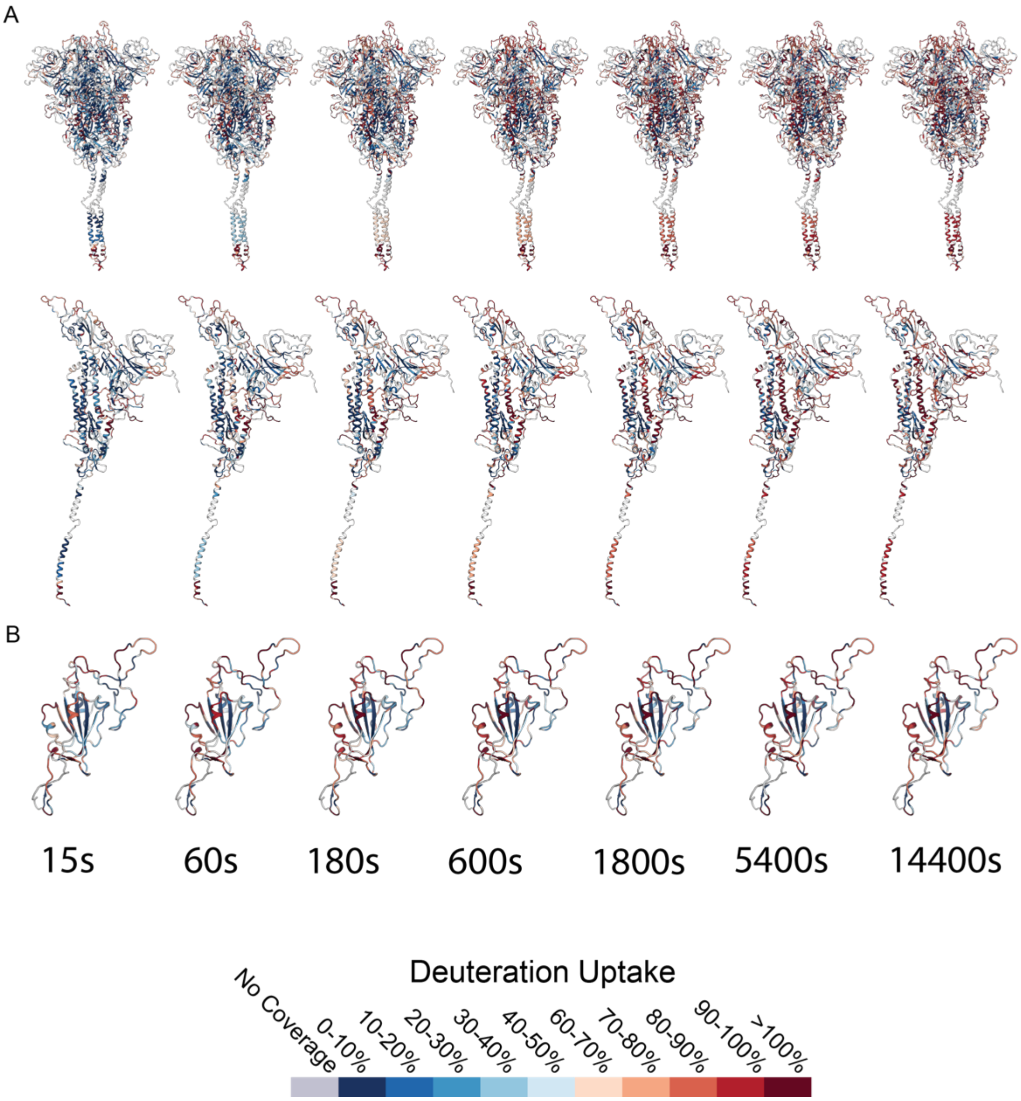


**Fig. S2. Spike HDX-MS results as a function of time**
**(A)** Deuterium uptake for each S-2P experimental time point mapped to the structure of the prefusion trimer and a single protomer (model from L. Casolino et al.). Per residue deuteration calculated from all peptide data by HDExaminer 3. **(B)** Deuterium uptake for each Apo-RBD experimental time point mapped to the structure of the RBD (single RBD from a full-length spike trimer model from L. Casolino et al.)

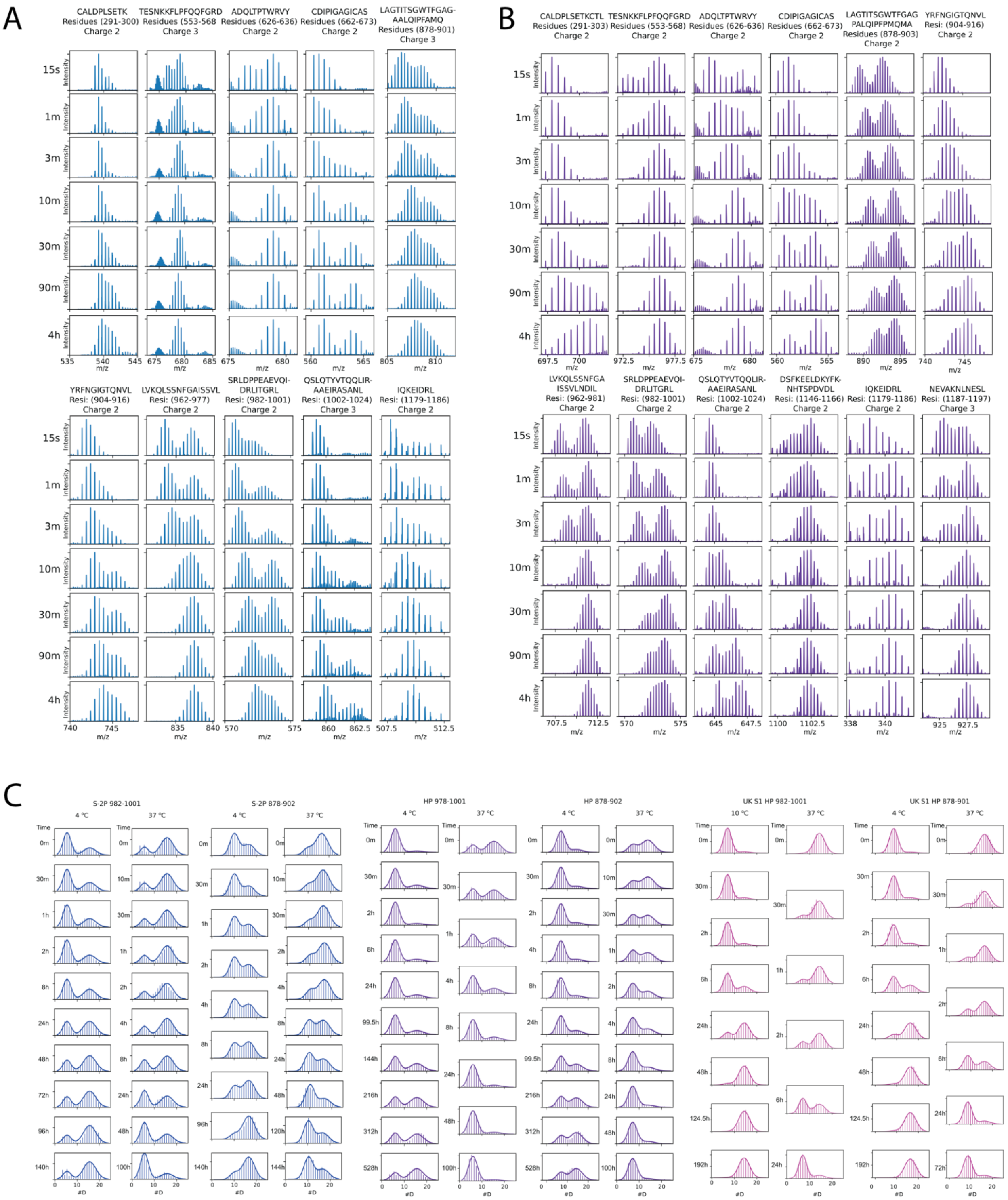


**Fig. S3. Bimodal peptide spectra observed in continuous and pulse-labeling HDX-MS experiments**

All peptides with observed bimodal behavior for both **(A)** S-2P and **(B)** HexaPro. **(C)** Bimodal peptides used to quantify the relative populations of state A and B are shown with the resulting gaussian fits overlaid.

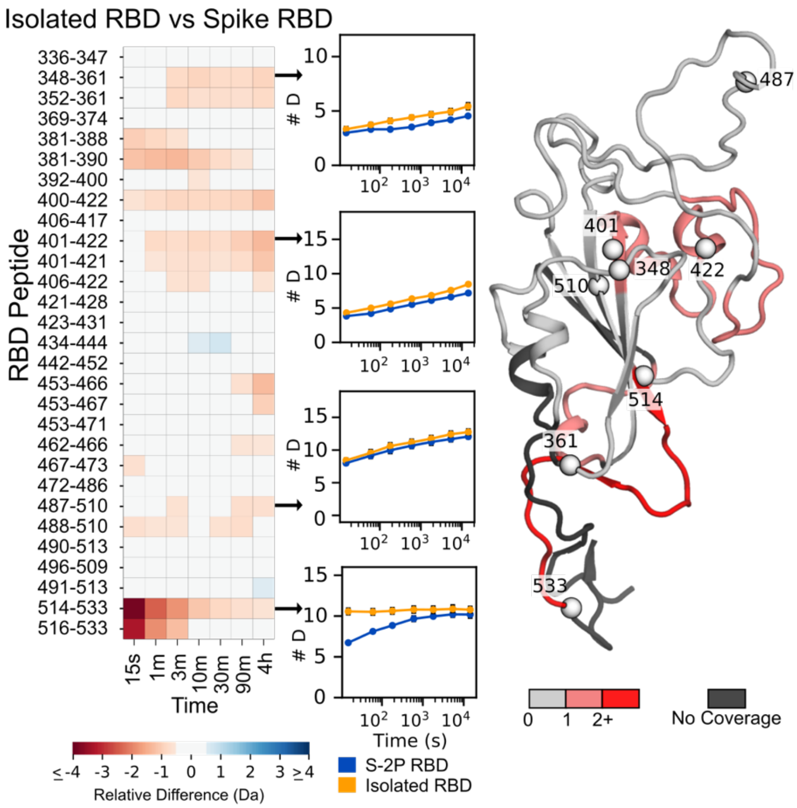


**Fig. S4. Comparison of HDX on RBD in isolation versus in S-2P**

Left: Heat map showing the difference in peptide deuteration for isolated RBD compared to the RBD in S-2P. Middle: Selected uptake plots of isolated RBD and S-2P RBD. Right: Structure of the RBD (model of a single RBD taken from a full-length spike trimer model from L. Casolino et al.) colored based on the maximum change shown in the heat map for that residue in any peptide. For reference, spheres are shown denoting the beginning and end of the peptides displayed in the uptake plots.

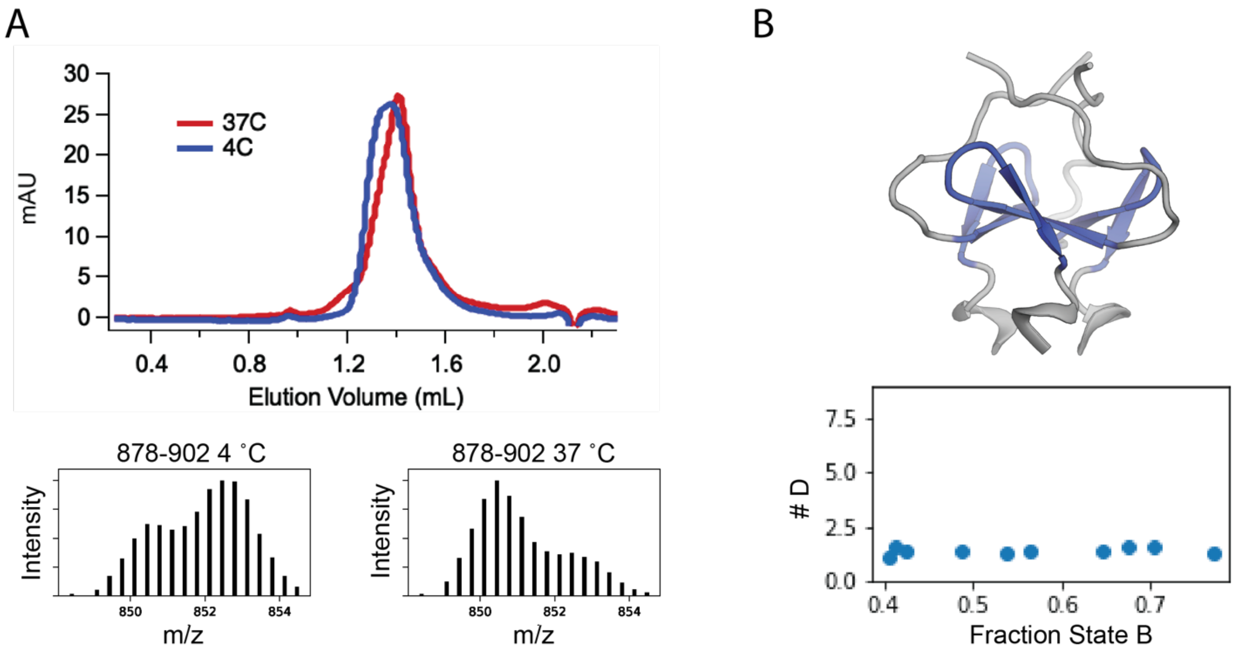


**Fig. S5. State A and state B are both Trimeric.**
**(A)** Top: SEC (Superose 6 increase 3.2/300) traces from S-2P after incubation at 37 °C and 4 °C. Bottom: MS spectra of a bimodal peptide (878 -902) from each sample taken immediately before SEC experiment. **(B)** Top: Structure of the T4 Fibritin trimerization domain with peptide shown in blue. Bottom: peptide deuterium uptake at one minute as a function of fraction state B
